# Screening of purine nucleoside analogues against intracellular *Toxoplasma gondii*

**DOI:** 10.64898/2026.02.11.705309

**Authors:** Hamza A. A. Elati, Serge Van Calenbergh, Lilach Sheiner, Harry P. de Koning

## Abstract

Toxoplasmosis remains a world-wide public health concern, especially for the immunocompromised. Although this population segment is increasing due to therapeutic interventions, organ transplants and infections including HIV, treatment relies almost exclusively on sulfadoxine and pyrimethamine, antifolates developed against malaria but with only moderate efficacy against acute toxoplasmosis and no effect on the chronic stage. Here we explore whether 7-substituted analogues of 7-deazaadenosine (tubercidin) that have shown remarkable efficacy against other protozoan pathogens, might also show anti-toxoplasmic activity. Tubercidin and a series of eleven 7-substituted analogues including 2’-deoxy and 3’-deoxyribofuranoses was tested against intracellular *Toxoplasma gondii* tachyzoites. The test compounds yielded EC_50_ values between 0.012 and 1.72 µM, well below those of the control drug sulfadiazine (11.9 µM) and the previously identified purine analogue adenosine arabinoside (Ara-A; 11.4 µM). The tubercidin analogues displayed at most moderate toxicity to HFF cells, with the most efficacious compound, 7-(3,4-di-Cl-phenyl)-3’-deoxytubercidin (FH8513) reaching a selectivity index of >2500. These nucleosides are most likely taken up by *T. gondii* through one of the four Equilibrative Nucleoside Transporters (ENTs) encoded by the parasites. However, deletion of *TgENT2* and/or *TgENT3* had no effect on the EC_50_ values, and deletion of *TgAT1* actually sensitised the tachyzoites to most of the tubercidin analogues. We propose that these nucleosides are internalised through the TgENT1 uridine transporter and that the sensitisation in ΔTgAT1 cells is the result of reduced uptake of adenosine that competes with the tubercidin analogues for metabolic enzymes such as adenosine kinase.

## 1. Introduction

*T. gondii* is an obligate intracellular parasitic protozoan of the phylum Apicomplexa that causes toxoplasmosis. It infects a wide range of hosts, including humans and all warm-blooded land and sea animals (Dubey, 1998; Su et al., 2003; Montoya and Liesenfeld, 2004). In humans, toxoplasmosis is a significant public health issue, affecting around one-third of individuals worldwide (Montoya and Liesenfeld, 2004; Molan et al., 2019). Infection can cause serious health problems, and even death in individuals with insufficient immune response, such as those with organ transplants, leukaemia, or AIDS. In such cases, the *T. gondii* parasite may reactivate from its chronic stage, in which it persists as tissue cysts containing bradyzoites. This reactivation can result in severe disease particularly affecting the central nervous system and the eye, with toxoplasmic encephalitis, focal neurological lesions and retinochoroiditis being the most common clinical manifestations (Luft and Remington, 1992; Bowen et al., 2016; Elsheikha et al., 2021). In pregnancy, the infection can be transmitted to the foetus and cause congenital toxoplasmosis, resulting in abortion, neonatal death, or congenital foetal abnormalities such as encephalitis, hydrocephalus, and retinochoroiditis that may appear at birth or later in life (Dunn et al., 1999; Montoya and Liesenfeld, 2004; Khan and Khan, 2018).

Although there has been some progress towards a toxoplasmosis vaccine, significant challenges remain (Zhang et al., 2023). Thus, currently, the principal therapeutic approach to contain toxoplasmosis involves chemotherapy to disrupt the parasite’s ability to synthesise folate with pyrimethamine and sulfadiazine, which inhibit dihydrofolate reductase (DHFR) and dihydropteroate synthase (DHPS), respectively; co-administration of folinic acid mitigates the side effects of these antifolates on the host (Alday and Doggett, 2017; Dunay et al., 2018). However, there is currently no therapy that is also effective against the chronic stage of the disease (Dunay et al., 2018), and new therapeutic strategies are urgently needed.

*T. gondii* is a purine auxotroph incapable of *de novo* purine biosynthesis and therefore relies on purine salvage from its host for its purine needs (Perrotto et al., 1971; Schwartzman and Pfefferkorn, 1982). This highlights the potential of the purine salvage system as an anti-toxoplasmic drug target and thus of purine analogues to treat toxoplasmosis. However, there have been few efforts to identify anti-toxoplasmic purine analogues, apart from extensive work on 6-benzylthioinosine and its analogues, by the group of Mahmoud el Kouni *(e*.*g*. (Al Safarjalani et al., 2010; Kim et al., 2010). Additionally, 6-thioxanthine and Ara-A were found to exert moderate parasitostatic activity against *T. gondii* tachyzoites (Pfefferkorn and Pfefferkorn, 1976; Pfefferkorn et al., 2001). However, 2’,3’-dideoxyinosine (didanosine, ddI), a drug developed to treat HIV, has shown activity against *T. gondii in vitro* and *in vivo*, including against tissue cysts (Sarciron et al., 1997; Sarciron et al., 1998; Gherardi et al., 2000). The latter observation shows that, unlike the first-line antifolates, some purine antimetabolites may actively destroy *T. gondii* bradyzoites as well as the rapidly replicating tachyzoites but this important observation has received relatively little attention. Indeed, very recently, it was reported that depletion of one of the *T. gondii* Equilibrative Nucleoside Transporter (ENT) genes, *TgENT1*, led to growth arrest (Messina et al., 2025; Qian et al., 2025; Elati et al., 2026) and that expression of two other ENTs, *TgAT1* and *TgENT3*, is necessary for bradyzoite differentiation (Messina et al., 2025).

Purine auxotrophic protozoa generally express multiple purine nucleoside and/or nucleobase transporters to efficiently acquire purines from the host environment (Landfear et al., 2004; Campagnaro and de Koning, 2020). The protozoan carriers commonly display far higher substrate affinity than their human counterparts (Wallace et al., 2002) and utilise the protonmotive force to actively accumulate substrates in the cell (De Koning and Jarvis, 1997; Stein et al., 2003). These qualities are exceedingly favourable to target purine antimetabolites into protozoa, achieving a high degree of selectivity over human cells in the translocation step (Luscher et al., 2007; Munday et al., 2015; Hulpia et al., 2019a). To date, the only nucleoside/nucleobase transporters identified in protozoa have been of the ENT family (Campagnaro and de Koning, 2020; Ungogo et al., 2023), and this is also true for *T. gondii* (Chiang et al., 1999; Messina et al., 2025; Qian et al., 2025; Elati et al., 2026). We very recently reported the involvement of TgENT1 in the uptake of not just uracil and uridine but also 5-fluoropyrimidines (Elati et al., 2026) – pyrimidine antimetabolites with higher *in vitro* anti-toxoplasmic activity than the first line antifolate sulfadiazine (Elati et al., 2023).

The above shows the potential for nucleoside and nucleobase antimetabolites against toxoplasmosis. Indeed, purine nucleosides and nucleobases have over the last few years emerged as serious drug candidates against a range of protozoan pathogens (Hulpia et al., 2019a; Bouton et al., 2021; Natto et al., 2021; Ilbeigi et al., 2025; Tashie et al., 2025). Therefore, the aim of this study was to investigate the activity of a series of purine nucleoside and nucleobase analogues against *T. gondii* RH tachyzoites strains *in vitro*. To gain insight in the role of ENT transporters, we also included strains in which ENT genes were deleted.

## 2. Methods

### 2.1. Cell lines and cultures

The F3 tomato strain (RH Δku80 TATi), derived from RH (Sheiner et al., 2011; Elati et al., 2023), in here referred to as RH, was used during the drug screening, alongside ΔTgAT1 (TGGT1_244440-KO), ΔTgENT2 (TGGT1_359630-KO), ΔTgENT3 (TGGT1_233130-KO), and DK (Δ(TgENT2/TgENT3)) cells that were also derived from RH (Elati et al., 2026). All *T. gondii* tachyzoites used in this project were cultured in human foreskin fibroblasts (HFF), sourced from ATCC (SCRC-1041). Parasites were passaged routinely as confluent monolayer of HFF in Dulbecco’s Modified Eagle’s Medium (DMEM) containing 4.5 gL^-1^ glucose, supplemented with 10% (v/v) foetal bovine serum, 4 mM L-glutamine and penicillin/streptomycin antibiotics, and grown at 37 °C with 5% CO_2_.

### 2.2. *Drug sensitivity assays for* Toxoplasma gondii

Drug sensitivity assays for *T. gondii* tachyzoites were assessed as previously described (Elati et al., 2023). Briefly, HFF were grown to confluence in black, clear bottom 96 well plates. Test compounds and positive control sulfadiazine were prepared in DMEM and serially diluted across the plate; the final column in the plate served as only infected cells. Freshly egressed tachyzoites (∼1000/well) were added and plates were incubated for 6 days at 37 °C with 5% CO_2_. Fluorescence was measured using a PHERAstar microplate reader (excitation: 540 nm; emission: 590 nm). EC_50_ values were obtained in GraphPad Prism 8.0 using a 4-parameter variable-slope sigmoid curve. All assays were performed in triplicate and repeated independently 3–5 times

### 2.3. Drug cytotoxicity assay for HFF cells

Cytotoxicity was as previously described (Elati et al., 2023), using phenylarsine oxide (PAO) as positive control. Plates were incubated for 6 days at 37 °C in a humidified atmosphere containing 5% CO_2_. On day 6,10 μL of resazurin solution (12.5 mg/100 mL ddH_2_O) was added to each well, including media-only wells to calculate background fluorescence. After incubation for 3–4h, fluorescence was measured using a PHERAstar plate reader (BMG Labtech) with excitation at 540 nm and emission at 590 nm. EC_50_ values were calculated in GraphPad Prism 8.0 by fitting a four-parameter sigmoidal dose-response curve with variable slope. All experiments were performed in triplicate and repeated 3–5 times.

### 2.4 Chemicals

Sulfadiazine, NBMPR, resazurin sodium salt, Tubercidin and Phenylarsine Oxide were bought from Sigma-Aldrich (Poole, UK). Pyrimethamine was from Fluka; Ara-A was from ICN Biomedicals. In-house nucleoside analogues TH1003; FH5319; TH1012; FH11711; FH8512; FH9576; FH8512; FH9576; FH8513; FH9581; FH9574; FH8494 were all described previously as referenced in the text.

## 3. Results and Discussion

### 3.1. *Validation of a new drug screening protocol for drugs against intracellular* T. gondii *tachyzoites*

Most published protocols for screening anti-toxoplasma compounds used the MTT assay, which is a colorimetric assay that relies on tetrazolium salt and is used for accessing cell viability, cell proliferation and cytotoxicity for the host cells (Choi and Lee, 2018). Here, we adapted a protocol that was used to evaluate antiretroviral compounds against *T. gondii* tachyzoites (Wang et al., 2019). We prepared serial dilutions of compounds in 96-well plates containing HFF monolayer cells and then added 1×10^4^ cells/mL of freshly lysed tachyzoites (5-fold fewer than Wang et al.), followed by incubation for 6 days at 37 °C and 5% CO_2_ in a humidified incubator. The lower parasite density allowed us to avoid the washing step, used by Wang et al. to remove the excess of extracellular tachyzoites; this adaptation should increase the assay’s reproducibility as it leaves the host cell monolayer undisturbed.

We tested our modified protocol using a set of standard compounds (sulfadiazine, pyrimethamine, Ara-A and 6-*S*-[(4-nitrophenyl)methyl]-6-thioinosine (NBMPR)) against *T. gondii* RH tachyzoites in 2 – 4 biological repeats (Figure 1), yielding highly consistent results. Ara-A has been shown to have moderate activity against *T. gondii* (Pfefferkorn and Pfefferkorn, 1976) and to be ineffective against tachyzoites lacking a functional TgAT1 nucleoside transporter (Chiang et al., 1999). Its modest activity was thought to be the result of relatively low affinity for the TgAT1/Tg244440 transporter (Campagnaro et al., 2022). Our drug screening assay protocol was consistent with these previous reports, as the drug Ara-A showed modest activity against *T. gondii* with an EC_50_ of 11.4 ± 1.8 µM (n=4) (Table 1).

**Figure 1.**
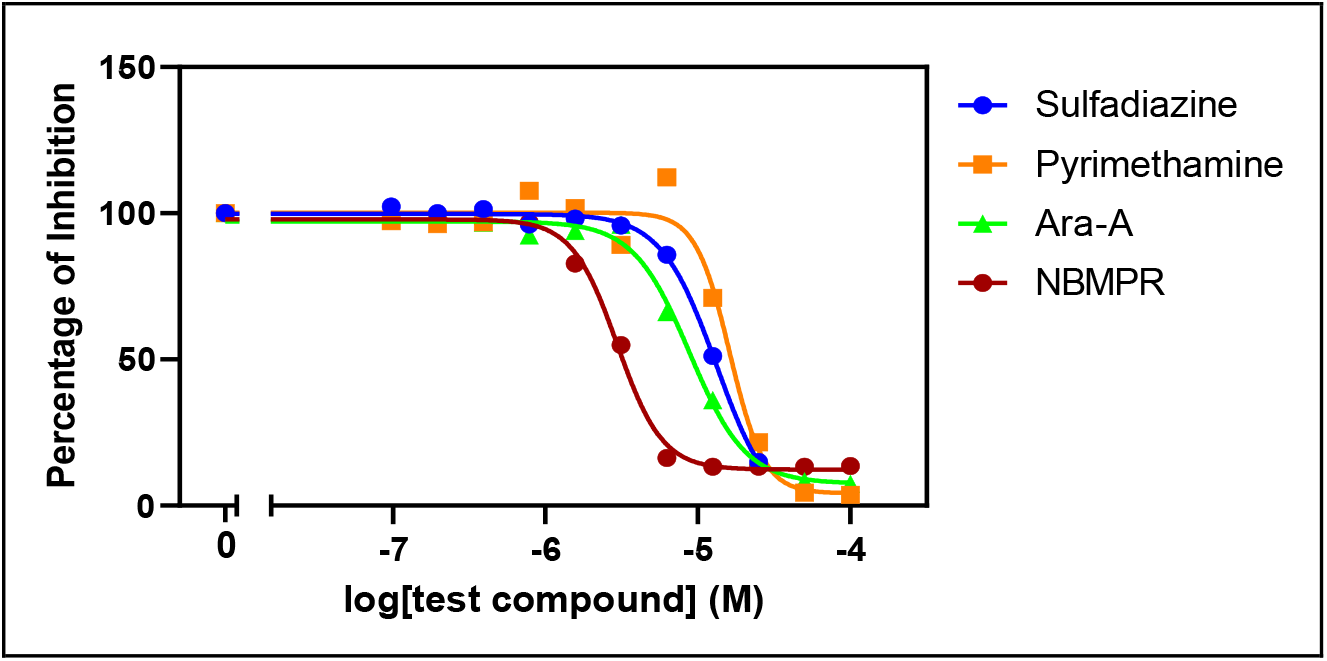
Drug sensitivity assay of *T. gondii* RH intracellular tachyzoites using standard treatments sulfadiazine and pyrimethamine, and the purine nucleoside analogues Ara-A and NBMPR. The figure shows a single representative experiment.

**Table 1:** In vitro drug screening and HFF cytotoxicity using purine nucleoside analogous on different cell lines of intracellular *T. gondii* RH tachyzoites, ΔTgAT1 and Δ(TgENT2/3).

| Compound ID | Modification<br>with respect to<br>tubercidin | RH control | ΔTgAT1 | | $\Delta$(TgENT2/3) | | HFF cytotoxicity | |
| --- | --- | --- | --- | --- | --- | --- | --- | --- |
| | | EC<sub>50</sub>± SEM<br>(μM) | EC<sub>50</sub>± SEM<br>(μM) | P value<sup>3</sup> | EC<sub>50</sub> ± SEM<br>(μM) | P value<sup>3</sup> | EC<sub>50</sub> ± SEM<br>(μM) | SI |
| ara-A<br>(vidarabine) | - | 11.4 ± 1.8 | 89.6 ± 9.23 | 0.0002 | 10.7 ± 1.9 | 0.80 | ND |  |
| NPMPR | - | 3.19 ± 0.14 | 2.06 ± 0.34 | 0.02 | 2.79 ± 0.07 | 0.05 | NE, 100 | >31.3 |
| tubercidin | - | 0.19 ± 0.05 | 0.17 ± 0.03 | 0.68 | 0.17 ± 0.01 | 0.80 | ND | -- |
| TH1003<sup>1</sup> | 7-Br | 0.10 ± 0.01 | 0.49 ± 0.02 | 0.000002 | 0.09 ± 0.01 | 0.65 | 1.94 ± 0.36 | 19.4 |
| FH5319 | 7-I | 1.48 ± 0.12 | 1.52 ± 0.09 | 0.76 | 1.24 ± 0.12 | 0.19 | >100 | >33.7 |
| TH1012<sup>2</sup> | 7-(4-Cl-Ph) | 0.50 ± 0.07 | 0.12 ± 0.02 | 0.001 | 0.33 ± 0.08 | 0.14 | >25 | >50 |
| FH11711<sup>2</sup> | 7-(4-Cl-Ph)<br>2'-deoxy | 0.55 ± 0.10 | 0.12 ± 0.05 | 0.005 | 0.55 ± 0.13 | 0.97 | >50 | >90 |
| **FH8512<sup>2</sup>** | 7-(4-Cl-Ph)<br>3'-deoxy | **0.21 ± 0.02** | 0.023 ±<br>0.008 | 0.0001 | 0.20 ± 0.04 | 0.75 | >50 | **>240** |
| FH9576<sup>2</sup> | 7-(4-CF<sub>3</sub>-Ph)<br>3'-deoxy | 0.57 ± 0.05 | 0.25 ± 0.06 | 0.005 | 0.48 ± 0.05 | 0.22 | NE, 50 | >87.7 |
| **FH8513<sup>1</sup>** | 7-(3,4-di-Cl-Ph)<br>3'-deoxy | **0.012 ±**<br>**0.002** | 0.0022 ±<br>0.0005 | 0.007 | 0.01 ± 0.002 | 0.91 | >25 | **>2500** |
| FH9581<sup>1,2</sup> | 7-(2,4-di-Cl-Ph)<br>3'-deoxy | 1.02 ± 0.20 | 0.28 ± 0.06 | 0.01 | 0.87 ± 0.11 | 0.53 | >25 | > 24.5 |
| FH9574<sup>2</sup> | 7-(4-*i*Pr-Ph)<br>3'-deoxy | 1.72 ± 0.14 | 1.15 ± 0.12 | 0.02 | 1.72 ± 0.13 | 0.97 | >100 | >30 |
| FH8494<sup>2</sup> | 4-(MeO-Ph)<br>3'-deoxy | 1.08 ± 0.14 | 0.45 ± 0.16 | 0.02 | 1.08 ± 0.01 | 0.96 | >25 | >23.2 |
| **FH9575<sup>1</sup>** | 7-(2-naphthyl)<br>3'-deoxy | **0.049 ±**<br>**0.007** | 0.011 ±<br>0.003 | 0.001 | 0.03 ± 0.005 | 0.07 | >10 | **>200** |
| Sulfadiazine |  | 11.9 ± 0.66 | 3.95 ± 0.29 | 0.0002 | 7.49 ± 0.28 | 0.002 | >100 | >84 |
| PAO | Phenylarsine<br>oxide | -- | -- | -- | -- | -- | 0.24 ± 0.03 | ND |

NBMPR and its 5′-monophosphate (NBMPR-P) reportedly exert selective toxicity to *T. gondii* with IC_50_ values of 10.2 µM and 9.9 µM, respectively, without apparent cytotoxicity on uninfected HFF monolayer cells up to 100 µM (El Kouni et al., 1999). Our testing of NBMPR against *T. gondii* RH tachyzoites was somewhat more sensitive and produced an EC_50_ of 3.19 ± 0.14 µM (n=4); there was no apparent cytotoxicity for NBMPR on uninfected HFF cells at concentrations up to 100 µM. (Table 1). Both purine analogues displayed a higher *in vitro* anti-toxoplasmic activity than either of the first-line antifolates (Figure 1).

### 3.2. The anti-toxoplasmic activity of AraA is dependent on uptake via TgAT1 but the activity of tubercidin is not

EC_50_ values were determined against *T. gondii* RH intracellular tachyzoites expressing the Tomato Red Fluorescent Protein, and on TgENT knockout cell lines ΔTgAT1, ΔTgENT2, ΔTgENT3, and DK (Δ(TgENT2/TgENT3)). Sulfadiazine was used as a positive control throughout for the screening against *T. gondii* cell lines and phenylarsine oxide (PAO) was used as a positive control for HFF cytotoxicity.

The sensitivity of intracellular ΔTgAT1 tachyzoites to Ara-A was strongly decreased (89.6 ± 9.23 µM, *P* < 0.0002) relative to *T. gondii* RH tachyzoites (Resistance Factor (RF) = 7.9). This is consistent with the resistance to Ara-A that was acquired after the loss of TgAT1 (Chiang et al., 1999), and the DK strain did not show any changes in Ara-A sensitivity relative to RH tachyzoites (EC_50_ value of 10.7 ± 1.9 µM (*P* > 0.05)). Tubercidin (7-deazaadenosine) was very active against RH tachyzoites with an EC_50_ value of 0.19 ± 0.05 µM (n=4), which was not significantly different against ΔTgAT1 and DK cells (*P* > 0.05).

### 3.3. Identification and SAR of tubercidin analogues with anti-toxoplasmic activity

A set of adenosine analogues was tested on *T. gondii* RH tachyzoites and different knockout strains of TgENT cell lines (Table 1). We selected part of an in-house library of 7-substituted analogues of tubercidin (7-deazaadenosine) as these have previously shown low toxicity to human cells combined with promising activity against a multiple protozoan pathogens, including *Trichomonas vaginalis* (Natto et al., 2021), *Trypanosoma cruzi* (Fiuza et al., 2022), *Trypanosoma congolense* (Mabille et al., 2022; Ungogo et al., 2023), *Leishmania infantum* (Lin et al., 2022), *Trypanosoma vivax* (Ilbeigi et al., 2025) and *Trypanosoma brucei* (Hulpia et al., 2019a; Hulpia et al., 2019b; Hulpia et al., 2020). While tubercidin has inherent anti-protozoal activity of its own (Iovannisci et al., 1984; Drew et al., 2003), it is too toxic to be a clinical candidate. Introduction of selected substitutions at position 7 reduce this toxicity (Hulpia et al., 2019a). The series is mainly comprised of 3’-deoxyribofuranose analogues as these were previously observed to be more potent against *T. cruzi* (Hulpia et al., 2018) and *T. b. brucei* and *T. b. rhodesiense* (Hulpia et al., 2019b) and here we show that they also proved beneficial for anti-toxoplasmic activity.

All analogues showed better activity than sulfadiazine, AraA and NBMPR against *T. gondii* RH tachyzoites, with EC_50_ values ranging from (0.012 to 3.19 µM) (Table 1; Figure 2). This indicated that 7-substituted tubercidin analogues are very well taken up by *T. gondii* and this was further investigated using the transporter knockout cell lines.

**Figure 2.**
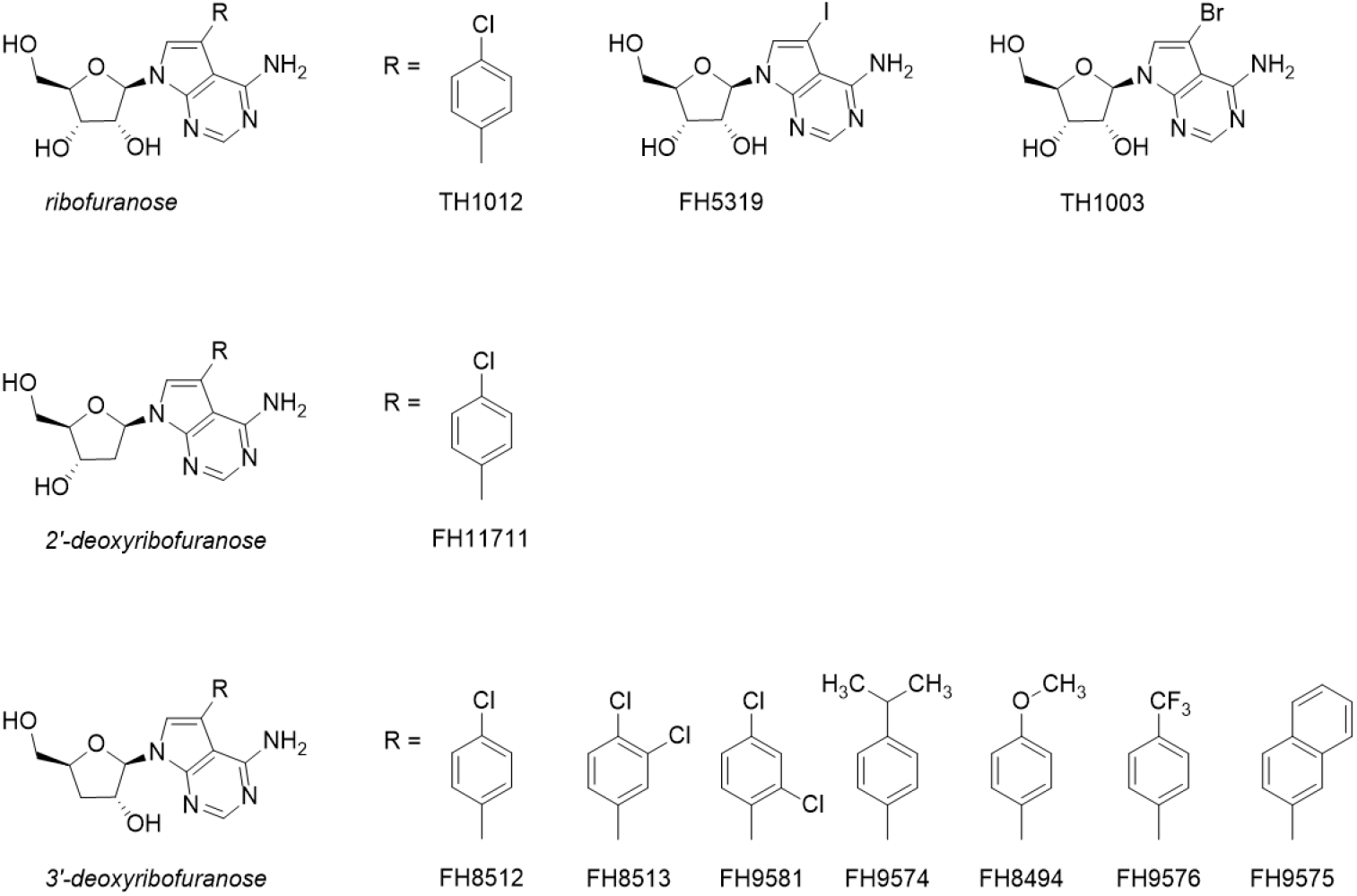
Structures of nucleoside analogues of 7-deaza-adenosine, 3’-deoxy-7-deaza-adenosine and 2’-deoxy-7-deaza-adenosine.

7-Bromotubercidin (TH1003) yielded a very promising EC_50_ value of 0.10 ± 0.01 µM, but showed a toxic effect against HFF cells with an EC_50_ ∼2 µM. The iodo analogue (FH5319) was less active (EC_50_ = 1.48 ± 0.12 µM (n=4)) but was devoid of cytotoxicity up to 100 µM. 7-(4-Chlorophenyl)tubercidin) (TH1012) was as active as tubercidin against RH tachyzoites with an EC_50_ of 0.50 ± 0.07 µM, and non-toxic to HFF cells up to at least 25 µM. This ribofuranose was earlier reported to have activity against intracellular *T. cruzi* amastigotes, with an EC_50_ of 0.47 µM (Hulpia et al., 2018), and particularly against *T. vaginalis*, for which it displayed an EC_50_ of 35 nM (Natto et al., 2021). In the latter study, no toxicity was observed against Human Embryonic Kidney (HEK293) cells or *in vivo* in a mouse model of trichomoniasis after 5 daily p.o. doses of 25 mg/kg body weight. TH1012 was also shown to be metabolically stable and to reduced parasite burden in a Chagas disease mouse model by 99.8% (equal to benznidazole, positive control) at 25 mg/ kg b.i.d. p.o. for 5 consecutive days (Hulpia et al., 2018).

Removal of the 2’-OH of TH1012 afforded FH11711, which showed an EC_50_ value of 0.55 ± 0.10 µM, similar to that of TH1012, implying that the 2’-OH group is not important for its anti-toxoplasma activity. Certainly, protozoan adenosine transporters do not require this hydroxyl for efficient uptake (Ogbunude et al., 1991; De Koning and Jarvis, 1999; De Koning et al., 2003). However, the 3’-deoxy-analogue of TH1012, *i*.*e*. FH8512, showed a > 2-fold better inhibitory effect on *T. gondii* RH tachyzoites (EC_50_ = 0.21 ± 0.02 µM (n=3)) and an EC_50_ >50 µM against HFF cells. It had also displayed a 10-fold stronger effect against *T. cruzi* amastigotes and was similarly effective in the mouse model as benznidazole and TH1012 (Hulpia et al., 2018). We therefore explored the series of 7-substituted 3’-deoxytubercidin analogues further.

The 3,4-dichlorophenyl analogue FH8513 (Bouton et al., 2021; Fiuza et al., 2022) emerged as the most potent analogue against intracellular tachyzoites with an EC_50_ value as low as 0.012 ± 0.002 µM (n=3). Notably, this compound displayed 1000-fold higher potency than sulfadiazine. Furthermore, it showed low cytotoxicity on HFF monolayers (EC_50_ > 25 µM), resulting in a selectivity index of more than 2500. Interestingly, the 2,4-dichlorophenyl isomer (FH9581) (Natto et al., 2021) was more than 80-fold less active, indicating that subtle structural modifications may have a very strong impact on the anti-toxoplasmic activity.

The 7-naphthyl-substituted analogue FH9575 showed much better activity against RH tachyzoites (EC_50_ = 0.049 ± 0.007 µM) than the 7-(4-iPr-phenyl) substituted analogue FH9574 (EC_50_ = 1.72 ± 0.14 µM) and 7-(4-CF_3_-phenyl) substituted analogue FH9576 (EC_50_ = 0.57 ± 0.05 µM) and also displayed good selectivity (SI > 200). FH9574 showed a cytostatic effect on HFF cells around 20 µM while its cytocidal effect became apparent at 100 µM. FH9576 displayed a slight cytostatic effect above 5 µM and a cytocidal effect above 50 µM. 7-(4-MeO-phenyl) substitution of 3’-deoxytubercidin (FH8494) resulted in significantly reduced activity compared to FH8512 and FH8513, suggesting that an electron-poor state of the aryl ring may favour activity against *T. gondii* tachyzoites.

### 3.4. Transporter dependence of anti-toxoplasmic activity of tubercidin analogues

The 7-substituted tubercidin analogues were all tested on the ΔTgENT2, ΔTgENT3 and DK (Δ(TgENT2/TgENT3)) cell lines but no significant differences with the control RH strain EC_50_ values were observed, so we completed triplicate determinations only with the DK strain and have not included the single experiments with ΔTgENT2 and ΔTgENT3 in Table 1. The result nevertheless clearly establishes that TgENT2 and TgENT3 are not determinative for the uptake of these antimetabolites.

Although ΔTgAT1 cells were identically sensitive to tubercidin and 7-iodotubercidin FH5319 as RH, this knockout strain was significantly less sensitive to the 7-Br analogue TH1003. More surprisingly, all 7-aryl substituted analogues showed significantly increased activity against ΔTgAT1 tachyzoites, by an average of 4.1-fold. This is not due to an upregulation of other TgENT transporters in the ΔTgAT1 strain (Elati et al., 2026) and, considering that this effect is specific to, and consistent for, the 7-aryl-3’-deoxytubercidin analogues seems to make it unlikely that it reflects upregulation of a relevant metabolic enzyme such as adenosine kinase (AK), considered to be the most important purine salvage pathway in *T. gondii* (Ngo et al., 2000; Chaudhary et al., 2004).

It is clear that tubercidin and its highly potent anti-toxoplasmic 7-aryl-3’-deoxyanalogues, such as FH8513, are not cross-resistant with AraA, and not dependent for uptake on TgAT1, TgENT2 or TgENT3 – which leaves TgENT1 as the only other known nucleoside transporter gene encoded by *T. gondii* (Messina et al., 2025; Qian et al., 2025; Elati et al., 2026). We have recently shown that TgENT1 is a high affinity uridine/uracil transporter with relatively low affinity for adenosine (Elati et al., 2026). While tubercidin and its analogues may be expected to enter through an adenosine transporter, as it does in *T. brucei* (De Koning and Jarvis, 1999; Geiser et al., 2005), in *T. cruzi* tubercidin enters through the TcrNT2 thymidine transporter (Aldfer et al., 2022a) and in *Leishmania spp* through the NT1 adenosine/uridine transporter (Vasudevan et al., 1998; Aldfer et al., 2022b). This shows a compatibility of uptake of pyrimidine nucleosides, adenosine and/or tubercidin by a by the same transporter, as also seen for human ENTs (Griffith and Jarvis, 1996).

We conclude, then, that it is highly likely that that the aryl-substituted tubercidin analogues are most likely taken up by tachyzoites through TgENT1, with the aryl substitutions possibly enhancing uptake through this carrier as is the case for the *T. brucei* P1 nucleoside transporter (Hulpia et al., 2019a; Hulpia et al., 2020). In this case, the enhanced efficacy of these compounds against the ΔTgAT1 strain can be explained in terms of adenosine uptake through TgAT1 (Chiang et al., 1999) competing with the purine antimetabolites for either a target or an activating enzyme like AK. Indeed, it is known from systematic structure-activity work by (Iltzsch et al., 1995) that 7-position substitutions greatly enhance the affinity of TgAK for tubercidin analogues and tubercidin is a substrate of the *Plasmodium chabaudi* AK (Schmidt et al., 1974). The deletion of the TgAT1 adenosine transporter reduces that competition, resulting in a modest increase in anti-toxoplasmic efficacy for the tubercidin analogues, following the same principle that sensitised *T. brucei* to 5-fluorouracil after deletion of a key enzyme in pyrimidine biosynthesis (Ali et al., 2013). It should be noted that the induction of resistance to the purine antimetabolite AraA resulted in the loss of its transporter, TgAT1 (Chiang et al., 1999). A similar resistance mechanism is much less likely for the tubercidin analogues as TgENT1 is an essential transporter (Messina et al., 2025; Qian et al., 2025), which could delay the onset of resistance to this class of purine antimetabolites.

## 4. Conclusion

*T. gondii* is a purine auxotroph (Krug et al., 1989; Chaudhary et al., 2004) that relies on the efficient salvage of large amounts of purines, principally adenosine through direct phosphorylation to nucleotides by AK, which is expressed at very high levels (Krug et al., 1989). Some 7-substituted tubercidin analogues have previously been shown to be high affinity substrates of TgAK (Iltzsch et al., 1995) and here we identify a series of 7-aryl-substituted 3’-deoxytubercidin analogues with up to 3 orders of magnitude stronger anti-toxoplasmic activity *in vitro* than sulfadiazine, a first-line drug, and low toxicity to HFF cells. These nucleosides are not dependent on uptake via TgAT1, TgENT2 or TgENT3 and we thus propose that they are substrates of the essential uridine carrier TgENT1, in line with tubercidin (analogues) sharing uridine transporters in other protozoa. We believe that the identified series should be further expanded on and that further studies should confirm the mechanism of action through TgENT1 and TgAK.

## Acknowledgements

This research was funded by a Wellcome Investigator Award (217173/Z/19/Z) and a Wellcome Discovery Award (310879/Z/24/Z)(to L.S.). HE was supported by a studentship from the government of Libya. S.vC. and H.dK. are members of the COST Action CA21111 “One Health drugs against parasitic vector borne diseases in Europe and beyond” and acknowledge the contributions of this EU-funded network to the development of antiparasite drugs. The authors would like to acknowledge Fabian Hulpia for the synthesis of the tubercidin analogues and Izet Karalic for excellent technical assistance.”

## Author Contributions

Conceptualization, H.P.dK.; methodology: L.S., H.A.A.E.; investigation, H.A.A.E.; formal analysis, H.A.A.E., H.P.dK.; resources: L.S., S.vC. writing—original draft preparation, H.P.dK.; writing—review and editing, H.A.A.E., S.vC., H.P.dK.; supervision, L.S., H.P.dK.; funding acquisition, L.S., H.P.dK.

